# Fluorescent reporter lines for auxin and cytokinin signalling in barley (*Hordeum vulgare*)

**DOI:** 10.1101/236018

**Authors:** Gwendolyn K. Kirschner, Yvonne Stahl, Jafargholi Imani, Maria von Korff Schmising, Rüdiger Simon

## Abstract

The phytohormones auxin and cytokinin influence the development and maintenance of plant stem cell niches. Although barley (*Hordeum vulgare*) is the fourth most abundant cereal crop plant, the knowledge about these important phytohormones in regard to the root and shoot stem cell niche in barley is still negligible. In this study, we analyse the influence of auxin and cytokinin on the barley root meristem and present reporter lines to describe the auxin and cytokinin signalling output. Application of high concentrations of auxin and cytokinin to barley seedlings had a negative influence on barley root and meristem growth. The expression of the cytokinin reporter *TCSn* revealed that cytokinin signalling mostly takes place in the stele cells proximal to the QC and in the differentiated root cap cells, but can additionally be activated in the root stem cell niche by cytokinin application. Analysing signalling targets of auxin showed that a homologue of AtPLT1, HvPLT1, is expressed in a similar way as AtPLT1 in *Arabidopsis*, in particular in the QC and the surrounding cells. Furthermore, a homologue of the auxin PIN transporters PIN1, HvPIN1, was expressed in the root and the shoot meristem and polarly localizes to the plasma membrane. Its expression is regulated by cytokinin and the intracellular localisation is affected by BFA. With this study, we provide a valuable tool set of fluorescent barley reporter lines for auxin and cytokinin.

## Introduction

The phytohormones auxin and cytokinin are crucial regulators in plant development, for example in embryogenesis, phyllotaxis, gravitropism and root and shoot formation (Reinhardt et al., 2003; Benková et al., 2003; Friml et al., 2003; Gordon et al., 2009; Marchant et al., 1999). Auxin controlled gene expression is transcriptionally regulated by AUXIN RESPONSE FACTORS (ARFs), which bind to short DNA sequences termed auxin responsive elements (AuxRes) in the promoter of target genes. At low auxin concentrations, the co-repressor TOPLESS represses auxin-regulated transcription by mediating the binding of AUXIN/ INDOLE-3-ACETIC ACID (Aux/IAA) proteins to ARFs. Perception of auxin by the TRANSPORT INHIBITOR RESPONSE 1/AUXIN SIGNALING F-BOX (TIR1/AFB) family proteins, subunits of an SCF E3-ligase protein complex, target the Aux/IAA proteins for degradation via the ubiquitin-proteasome pathway, thereby leading to the activation of the ARFs and hence auxin responsive gene expression (reviewed in Saini et al., 2013). Cytokinins are perceived by histidine kinase receptors (AHKs) which carry an extracellular CHASE domain for hormone sensing. Cytokinin perception leads to autophosphorylation of the receptor kinase domain and subsequent transfer of the phosphoryl group onto a histidine phosphotransfer-protein (AHP). This enables AHP allocation to the nucleus and relay of the phosphoryl group to type-B response regulators (type-B ARRs), which in turn regulate transcription of cytokinin responsive genes. Among their targets are type-A ARRs which negatively influence cytokinin signalling, thereby creating a negative feedback loop (reviewed in Bishopp et al., 2011). Phytohormone distributions in a tissue can be visualized by the expression of reporter genes under the control of known auxin- or cytokinin-responsive elements, such as *DR5* and *DR5v2* for auxin signalling, and *Two Component signalling Sensor* (*TCS*) and *TCSnew* (*TCSn*) for cytokinin (Ulmasov et al., 1997; Liao et al., 2015; Zurcher et al., 2013). A more direct way to analyse the presence of auxin is monitoring the degradation of reporter proteins fused to the highly conserved SCF-TIR1 complex recognition domain (DII) of Aux/IAA proteins (Brunoud et al., 2012; Liao et al., 2015). In the *Arabidopsis* shoot apical meristem (SAM), these reporters uncovered auxin maxima in the primordia, and high levels of cytokinins in the center of the meristem and the primordia. In the root apical meristems (RAM), an auxin maximum is formed in the Quiescent Center (QC), in the columella initials and in differentiated columella cells, while cytokinin maxima are observed in the differentiated columella and the stele (Aida et al., 2004; Sabatini et al., 1999; Yang et al., 2017; Benková et al., 2003; Zurcher et al., 2013). Post-embryonic development of plant organs depends on the activity of meristems, and this specific phytohormone distribution was shown to be required for meristem patterning (Sabatini et al., 1999; Blilou et al., 2005; Reinhardt et al., 2003; Stahl and Simon, 2005). At least for auxin, fine-tuned short-distance transport is essential to establish and maintain its distribution pattern. PINFORMED (PIN) proteins serve as auxin efflux carriers and establish a directional auxin flow to maintain the auxin maximum (Wang et al., 2009a; Miyashita et al., 2010; Xu et al., 2005; Blilou et al., 2005; Carraro et al., 2006; Gallavotti et al., 2008; Reinhardt et al., 2003). Different PINs are expressed in specific, partially overlapping domains within the root meristem and polarly localize to the cell membrane, thereby exporting auxin only in one specific direction (Blilou et al., 2005; Carraro et al., 2006). Among downstream targets of auxin signalling in *Arabidopsis* are the *PLETHORA* (*PLT*) genes, which are members of the AINTEGUMENTA-like (AIL) subclass of the APETALA2/ethylene-responsive element binding proteins (AP2/EREBP) family of transcription factors (Aida et al., 2004). *AtPLT1* and *AtPLT2* are redundantly required for the embryonic specification of the QC and for the maintenance of root stem cells in *Arabidopsis* (Aida et al., 2004). The *AtPLTs* are expressed in the stem cell niche forming a concentration gradient with a maximum in the QC and the distal stem cells (DSCs), therefore mirroring auxin distribution (Aida et al., 2004; Galinha et al., 2007; Mähönen et al., 2014). Auxin and cytokinin signalling are connected at multiple points: for example, auxin induces ARR expression (Müller and Sheen, 2008; Moubayidin et al., 2013), while ARR1 promotes expression of the Aux/IAA gene *SHORT HYPOCOTYL2* (*SHY2*). Furthermore, cytokinin influences auxin transport by regulating the expression of auxin influx (LIKE AUXIN RESISTANT 2) and efflux carriers (PINs), causing a relocation of auxin (Dello Ioio et al., 2008; Ruzicka et al., 2009; Zhang et al., 2013). Homologues of PIN auxin efflux carriers, as well as auxin-responsive homologues of PLTs were also identified in monocots, indicating that the principles of auxin and cytokinin transport and signalling is conserved between monocots and dicots (Zhang et al., 2014; Li and Xue, 2011; Wang et al., 2009a; Xu et al., 2005; Carraro et al., 2006; Gallavotti et al., 2008; O’Connor et al., 2014). Even though barley (*Hordeum vulgare*) is the fourth most produced crop plant in the world (FAO statistics 2014; http://faostat.fao.org), highly salt tolerant in comparison to other cereal crops (Maas and Hoffman, 1977) and therefore a valuable model plant in regard to abiotic stresses, only very few studies about auxin and cytokinin exist for barley (Tagliani et al., 1986; Zalewski et al., 2014, 2010; Pospíšilová et al., 2016). Initial studies indicate a connection between cytokinin signalling and drought stress resistance in barley, which so far has not been fully explored (Pospíšilová et al., 2016). In this study, we analysed cytokinin signalling and downstream targets of auxin and cytokinin in the barley shoot and root meristem, utilizing phytohormone treatment, RNA *in situ* hybridisations and transgenic fluorescent reporter lines. Application of the hormones to barley seedlings impairs root growth and meristem maintenance. The expression pattern of the cytokinin signalling reporter *TCSn* reveals cytokinin signalling in the stele proximal to the QC and in the differentiated root cap cells. The homologue of the auxin-responsive gene *AtPLT1*, *HvPLT1*, is expressed in a pattern similar to *AtPLT1* in *Arabidopsis*, in particular in and around the QC. Furthermore, the putative auxin transporter HvPIN1a is expressed and polarly localised in the root meristem, its expression is regulated by cytokinin and the intracellular localisation is affected by BFA, similar to *Arabidopsis*. With our study we provide the first fluorescent reporter lines for phytohormone transport, signalling or responses in barley which serve as valuable tools to analyse the role of auxin and cytokinin in barley development.

## Material and Methods

### Plant growth

To monitor root growth and expression of reporter genes in the root, seedlings were grown on square plates as described in (Kirschner et al., 2017). For all experiments either the cultivar (cv.) Morex or Golden Promise were used as indicated. SAMs were monitored in plants grown 8 DAG on agar plates (Waddington stage I, “transition apex”) or plants grown on soil under greenhouse conditions for around 3 weeks (Waddington stage II “double-ridge”) (Waddington et al., 1983).

### Cloning

The *HvpPLT1:HvPLT1-mVENUS* construct was built by PCR amplification of a 1929 bp fragment upstream of the start codon of *HvPLT1* (MLOC_76811.2 on morex_contig_73008/ HORVU2Hr1G112280.5 (Mayer et al., 2012; Mascher et al., 2017)) as the putative promoter region from Morex genomic DNA (gDNA) and cloned by restriction and ligation via a *Asc*I site into a modified pMDC99 (Curtis and Grossniklaus, 2003). The whole *HvPLT1* coding region without stop codon (3433 bp) was amplified from Morex gDNA and inserted downstream of the promoter in the pMDC99 vector by Gateway cloning (Invitrogen). A C-terminal *mVENUS* (Koushik et al., 2006) was integrated downstream of the gateway site by restriction and ligation via *Pac*I and *Spe*I. The *HvpPIN1:HvPIN1-mV* construct was produced the same way, using 3453 bp upstream of the start codon of *HvPIN1* (AK357068/MLOC_64867 on morex_contig_101983/ HORVU6Hr1G076110.1 (Mayer et al., 2012; Mascher et al., 2017)) as putative promoter region and the whole *HvPIN1* coding region including the stop codon. The *mVENUS* sequence was inserted by restriction and ligation via a *Sma*I restriction site into the sequence coding for the central hydrophilic region of the HvPIN1 protein, as described for a PIN1 reporter construct in *Arabidopsis* (Benková et al., 2003). The insertion of mVENUS is depicted in Supplementary figure 8B. For the *TCSn:VENUS-H2B* cytokinin reporter construct, the *TCSn* regulatory sequence (Zurcher et al., 2013) was obtained in the pDONR221 gateway vector from Invitrogen and subsequently inserted by Gateway cloning into the modified pMDC99 vector. The auxin reporter construct *DR5v2:VENUS-H2B* was built by amplifying the *DR5v2* promoter from the *pGIIK/DR5v2::NLS-tdTomato* plasmid (kind gift of Dolf Weijers, (Liao et al., 2015)) and inserted by Gateway cloning into the modified pMDC99 vector. The pMDC99 modified for *TCSn:VENUS-H2B* and *DR5v2:VENUS-H2B* contained the gateway cassette, the coding sequence of *VENUS* (Nagai et al., 2002) and a T3A terminator, which were inserted by restriction and ligation with *Asc*I and *Sac*I from pAB114 (described in Bleckmann et al., 2010). Furthermore, it contains the coding sequence of *Arabidopsis HISTONE H2B* (AT5G22880) at the C terminus of the *VENUS* gene, inserted via restriction and ligation at a *Pac*I restriction site. The *DR5:ER-GFP* contains the auxin-response promoter *DR5* that consists of 9 inverted repeats of the 11 b-sequence 5’-CCTTTTGTCTC-3’, a 46-bp CaMV35S minimal promoter element, and a tobacco mosaic leader sequence as translational enhancer fused to endoplasmatic reticulum - targeted GFP (Benková et al., 2003; Friml et al., 2002). The plasmid was a kind gift from the Benková lab.

### Barley transformation

The barley cv. Golden Promise was used for transformation as described previously (Imani et al., 2011), tested for hygromycin resistance by growth on medium containing hygromycin and via PCR to detect the hygromycin gene. For root expression analysis, the seeds of the plants recovered from the transformed scutella were used (T1) and again tested for the presence of the reporter construct by PCR with primers binding in the gene of interest and the downstream reporter gene.

### Preparation of the reporter line samples

Clearing of the transgenic reporter lines was performed as described for pea root nodules with an altered fixation step (Warner et al., 2014). Root samples were fixed with 4 % para-formaldehyde in phosphate buffered saline (PBS) for 1 h with applied vacuum. Samples were incubated in the clearing solution for 1 week in darkness at 4 °C. The roots of plant lines with weak expression, or to be examined uncleared, were embedded in warm liquid 5 % (w/v) agarose in dH_2_O for stabilization and sectioned longitudinally in the center by hand with a razor blade. SAM preparation was carried out by removing all leaves from the SAMs and the expression was directly monitored without clearing.

### Cell wall and starch staining

Modified pseudo-Schiff propidium iodide (mPS-PI) staining and microscopy of the stained samples was performed as described previously (Kirschner et al., 2017).

### Treatments

The cytokinins 6-benzylaminopurine (6-BA) (Duchefa) and *trans*-zeatin (t-Z) (Sigma-Aldrich) were used, as well as the auxins NAA (Duchefa) and 2,4D (Duchefa). For phytohormone treatments of wild type plants, the hormones were added to the growth medium at the concentrations indicated in the results section. The mock control was treated with water. For phytohormone treatment of the *TCSn:VENUS-H2B* and *HvpPIN1:HvPIN1-mVENUS* and *DR5v2:VENUS-H2B* reporter lines, the phytohormones were added to PBS and the plates with 7 day-old seedlings were flooded with the hormone solution or pure PBS as mock control and incubated for 2 −3 h to allow phytohormone uptake. After removing the buffer, plates were placed back into the phytochamber at a 45 ° angle and examined 24 h later. Brefeldin-A (BFA) treatment of the *HvpPIN1:HvPIN1-mVENUS* reporter line was performed as described (Geldner et al., 2001). Roots were cut around 1 cm above the tip, which was then placed in PBS as mock control or PBS containing 50 μM BFA. Pictures of the outer cortex cell layers of the roots and the epidermis of the SAMs were taken at the time of treatment (0 h) and 2 h later.

### RNA *in situ* hybridisations

Probes for the *HvPLT1* mRNA were prepared from genomic DNA of the barley cv. Morex from the *HvPLT1* start to stop codon (3433 bp). The DNA was cloned into the pGGC000 entry vector of the GreenGate cloning system (Lampropoulos et al., 2013) and then amplified including the T7 and SP6 promoter sites by PCR. RNA probes were produced as described (Hejátko et al., 2006). The RNA probes were hydrolysed by adding 50 μl carbonate buffer (0.08 M NaHCO_3_, 0.12 M Na_2_CO_3_) to 50 μl RNA probe and incubation at 60 °C for 58 min. On ice, 10 μl 10 % acetic acid, 12 μl sodium acetate and 312 μl EtOH were added, the RNA was precipitated and dissolved in RNase-free dH_2_O. RNA *in situ* hybridisations were performed on roots of plants 8 days after germination (DAG) as described previously (Kirschner et al., 2017). Polyvinyl alcohol was added to a final concentration of 10 % to the NBT/BCIP staining buffer. Permanent specimens were created by washing the slides in 50 % EtOH, 70 % EtOH, 95 % EtOH and 100 % EtOH for 2 min each and for 10 s in xylol, and after drying, a few drops of Entellan (Merck) and a cover slip were added.

### Microscopy

The transgenic reporter lines with mVENUS or VENUS fluorophores were examined with a 40x water objective with a numeric aperture (NA) of 1.20 using the Zeiss confocal laser scanning microscope (LSM) 780. Yellow fluorescence was excited using a 514 nm Argon laser and the emission was detected between 519 and 620 nm. The pinhole was set to 2,24 airy units. Transmitted light pictures were recorded with a transmitted light detector (T-PMT). Pictures were recorded with the tile scan function with 10 % overlap, a threshold of 0.70 and automatically stitched using the microscope software. RNA *in situ* hybridizations were examined using a plan-neofluar 20x objective with a NA of 0.50 or a plan-neofluar 40x objective with a NA of 0.75 using the Zeiss Axioskop light microscope.

### Analysis

Picture analyses were carried out using Fiji (Schindelin et al., 2012). For root length measurements, the mean root length of all roots from a single plant were measured. For meristem length measurements, the border between meristem and elongation zone was defined by the first cell in the outermost cortex cell layer that doubled in cell length compared to its distal neighbour and analysis was carried out qualitatively from direct observation (as described in Dello Ioio et al. 2007). For analysing the DSC layers the starch-free cells of three columns in the center of the root cap below the QC were counted and the mean for one column was calculated. Only roots with mPS-PI stained starch-containing cells were used for analysis. For information about creation of the phylogenetic trees see Supplementary figure 6 and Supplementary figure 8A. The transmembrane domains of the PIN proteins were predicted using the TMHMM method (Krogh et al., 2001). Plots and statistics were created in R (R Core Team, 2015). Significance was determined by a two-tailed Student’s T-Test with the given p value. For image processing, Adobe Photoshop was used. Contrast and brightness were adjusted in the mPS-PI sample pictures manually to increase the cell wall and starch visibility. When the fluorescence brightness was compared, the same changes were performed equally for all samples. The surface of the SAMs was extracted in MorphoGraphX (de Reuille et al., 2015).

## Results

### Cytokinin inhibits root growth and root meristem maintenance

The effects of auxin and cytokinin signalling can easily be studied by manipulation of the hormone levels in the plant, for instance by externally adding an excess of the hormone, or by inhibiting biosynthesis or signal perception. Reduction of cytokinin levels by overexpression of degradation enzymes leads to enhanced root growth and longer meristems in *Arabidopsis* (Werner et al., 2010), whereas application of cytokinin reduces the root and meristem length (Ruzicka et al., 2009; Dello Ioio et al., 2007). While the effect of a manipulation of the cytokinin signalling and biosynthesis pathway was already studied in barley roots on a whole-organ level, it has not yet been analysed on a cellular level (Zalewski et al., 2010; Pospíšilová et al., 2016). To test the effect of cytokinin, we used both *trans*-zeatin (t-Z), a naturally occurring isoprenoid-type cytokinin (Podlešáková et al., 2012), and the synthetic cytokinin 6-benzylaminopurine (6-BA), both of which were shown to affect root and meristem length upon application in *Arabidopsis* (Ruzicka et al., 2009; Dello Ioio et al., 2007). 6-BA had a negative effect on root length in both used concentrations (1 μM and 10 μM) measured 10 DAG, while t-Z did not affect root growth significantly (Figure 1A). Root longitudinal growth can primarily be attributed to cell divisions in the meristematic zone and cell elongation in the elongation zone. Interestingly, the effect on the root meristems was much stronger for both, 6-BA and t-Z, and resulted in a reduction in meristem size (Figure 1B, C). Both hormones seem to affect meristem size by changes in cell division and/or differentiation rate, since a reduced cell number was responsible for the difference in overall meristem size, rather than the mean length of the meristematic cells (Supplementary figure 1A). Furthermore, both cytokinins reduced the width of the root meristem (Supplementary figure 2B). In summary, cytokinin application causes a reduction in root growth in barley which is due to a reduced meristem size and thereby a lower production of new root cells for growth in length or diameter.

**Figure 1:**
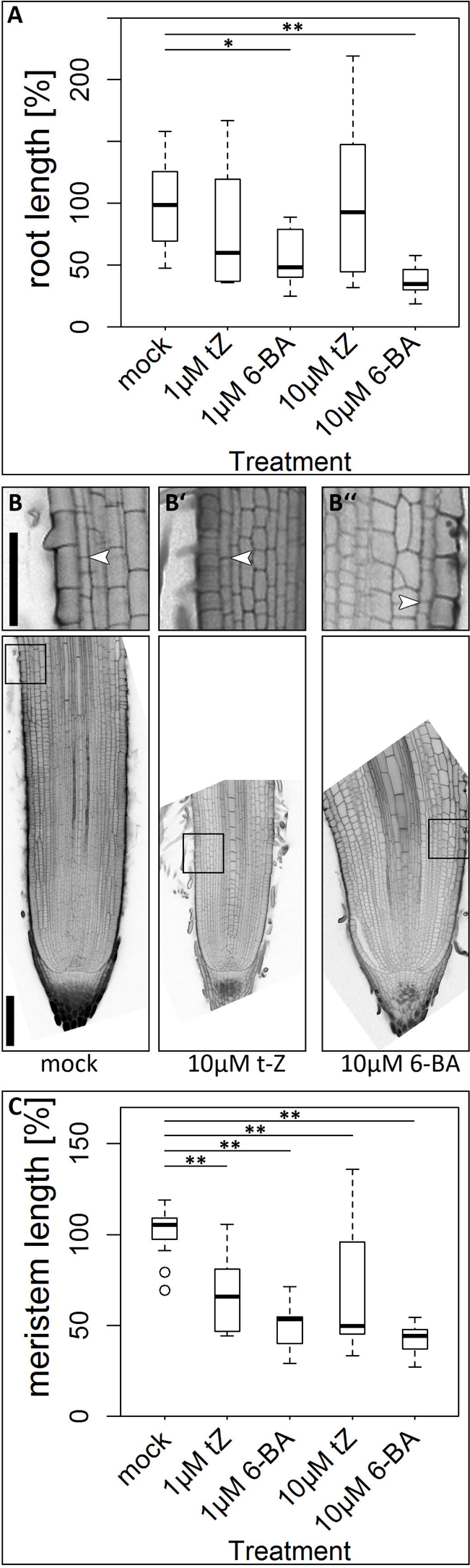
Root length and meristem size of the barley cv. Morex upon cytokinin treatment for 10 days. **A)** Root length after 10 day-treatment with cytokinin; experiment was performed twice; values normalized to mock-treated plants; n = 7-18 plants per data point. **B)-B’’)** Representative pictures of meristem phenotypes upon cytokinin treatment according to the captions; arrowheads mark the transition zones in the outer cortex layer; insets show magnifications of the transition zones; scale bars 200 μm (overviews) and 100 μm (magnified insets). **C)** Meristem size after 10-day cytokinin treatment, measured by meristem length; experiment was performed twice; n = 11-16 roots per data point; significance was determined using the two-tailed Student’s t test, * = p<0.05, **= p<0.001.

### The cytokinin signaling reporter *TCSn* is expressed in the stele and root cap and can be activated by cytokinin application

Since cytokinin application had an effect on root length and meristem size, we aimed to reveal the distribution of the phytohormone in the root. Therefore, we transformed barley cv. Golden Promise plants with a construct carrying a fluorescent *VENUS* reporter gene driven by the *TCSn* promoter within the vector backbone of pMDC99 (Curtis and Grossniklaus, 2003). This promoter carries concatemeric binding motifs for type-B ARRs combined with a minimal promoter, and responds to cytokinin signalling in *Arabidopsis* and maize protoplasts (Zurcher et al., 2013). In mature barley root apical meristems (8 DAG), we observed expression of the cytokinin reporter *TCSn:VENUS-H2B* in the differentiated root cap and the stele (Figure 2A’), but not in the metaxylem (Figure 2A’, white arrow head), the QC or the surrounding initials (Figure 2A’, gray arrow head), resembling the expression pattern of the *TCSn* promoter in *Arabidopsis* (Zurcher et al., 2013). 24 h treatment with 6-BA significantly increased *TCSn:VENUS-H2B* expression in the stele, while there was only a weaker response to t-Z (Figure 2B, C). Cytokinins activated *TCSn* expression also in the DSCs, the cortex and endodermis initials, the epidermis initials and in a layer of the QC adjacent to the root cap (Figure 2D). In control plants, expression in the DSCs, the QC or the surrounding initials could never be observed (Figure 2D). However, we could not observe expression of the *TCSn:VENUS-H2B* in the SAM of any of the transgenic lines (Supplementary figure 3).

**Figure 2:**
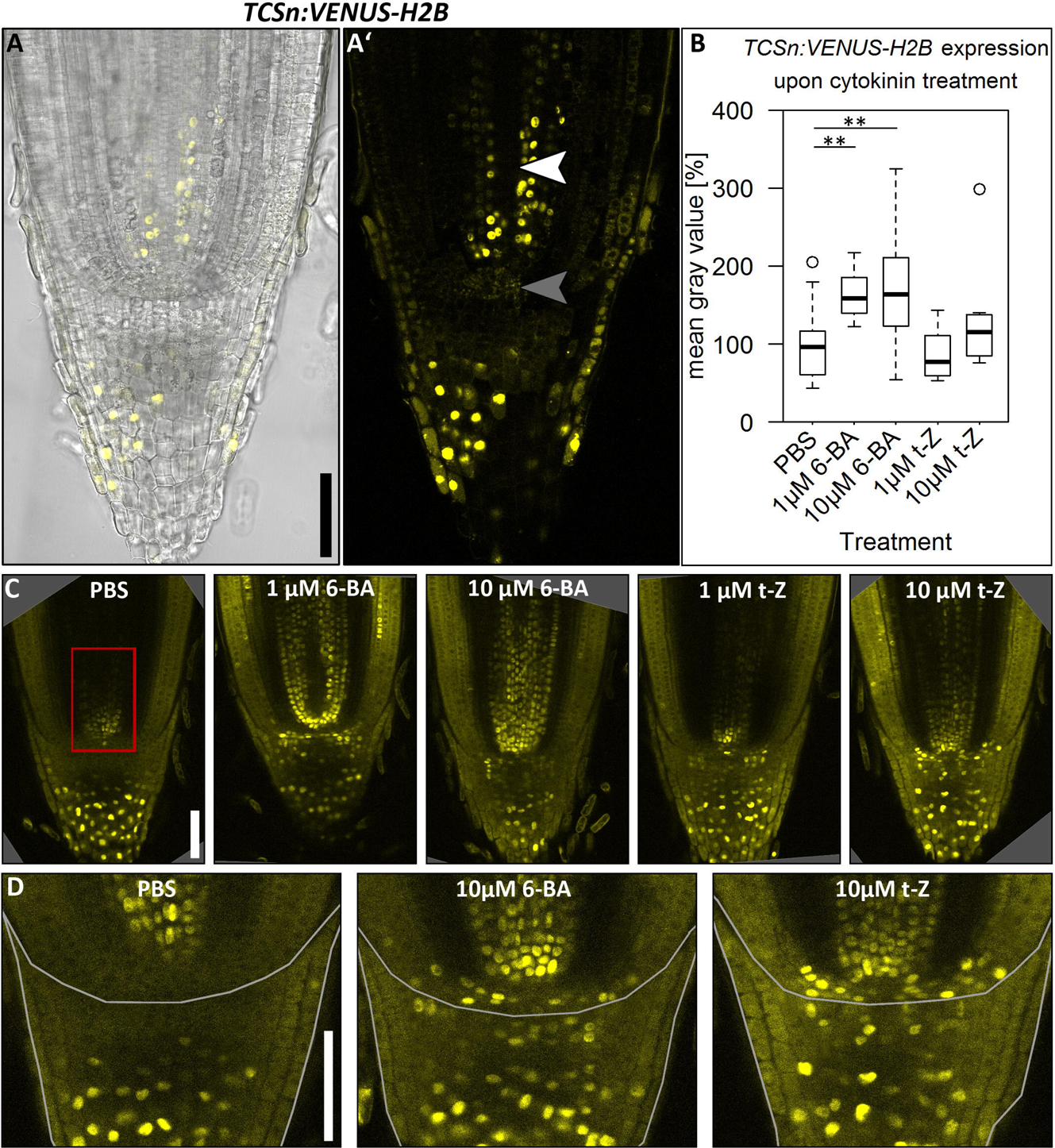
Expression of the cytokinin reporter *TCSn:VENUS-H2B* in the root meristem of the barley cv. Golden Promise 8 DAG. **A), A’)** *TCSn:VENUS-H2B* expression in untreated roots; transmitted light and VENUS emission (A)) and VENUS emission only (A’)); white arrow head in A’: metaxylem, gray arrow head in A’: QC; seven independent transgenic lines were examined and exhibit a similar expression pattern. **B)** Quantification of *TCSn:VENUS-H2B* expression by the mean gray value of the region marked by red box in C); mean gray value normalized to the PBS control; significance was determined using the two-tailed Student’s t test, ** = p<0.001. **C)** Representative pictures of *TCSn:VENUS-H2B* expression in root meristems upon 24 h of cytokinin treatment according to the captions; PBS without hormone was used as control; three independent transgenic lines were examined; the experiment was performed three times; n = 8-31 per treatment. **D)** Magnification of the stem cell niche and root cap of roots upon treatments indicated by the captions; expression in the cortex/ endodermis initials, the DSCs, the QC layer adjacent to the root cap and the epidermis initials (PBS: 0/21 roots, 1 μM 6-BA: 1/9 roots, 10 μM 6-BA 8/18 roots, 1 μM t-Z 2/9 roots, 10 μM t-Z 5/8 roots); the root cap border is marked with a white frame; for a better comparison between samples, roots were cleared before microscopy (C), D), E)); scale bars 100 μm.

### High auxin concentrations decrease root growth and negatively affect root meristem size

External application of auxin affects the root architecture of plants in regard to root length, meristem size and structure (Martínez-de la Cruz et al., 2015; Ruzicka et al., 2009; Carraro et al., 2006). In barley, root growth was inhibited by all auxins tested (Tagliani et al., 1986). To study barley root and meristem growth in more detail, we used the synthetic auxin NAA and the auxin analogue 2,4D, which cannot be exported from cells via auxin efflux carriers (Delbarre et al., 1996). Low (10 nM) and high concentrations (1 μM and 10 μM) were used in comparison, as auxins are known to have opposite effects on meristem size at different concentrations (Ruzicka et al., 2009). Growing barley plants on medium containing either no phytohormone or the different auxins for 10 days revealed that neither root length nor root meristem length are affected by low concentrations (10 nM) of NAA or 2,4D, but both are decreased at high auxin concentrations (1μM 2,4D, 1μM and 10μM NAA) (Figure 3A, B, C). The reduction in meristem length went together with a reduction in meristematic cortex cell number (Supplementary figure 1B). Additionally, treatments with high concentrations (1 μM and 10 μM) increased meristem width (Supplementary figure 2).

**Figure 3:**
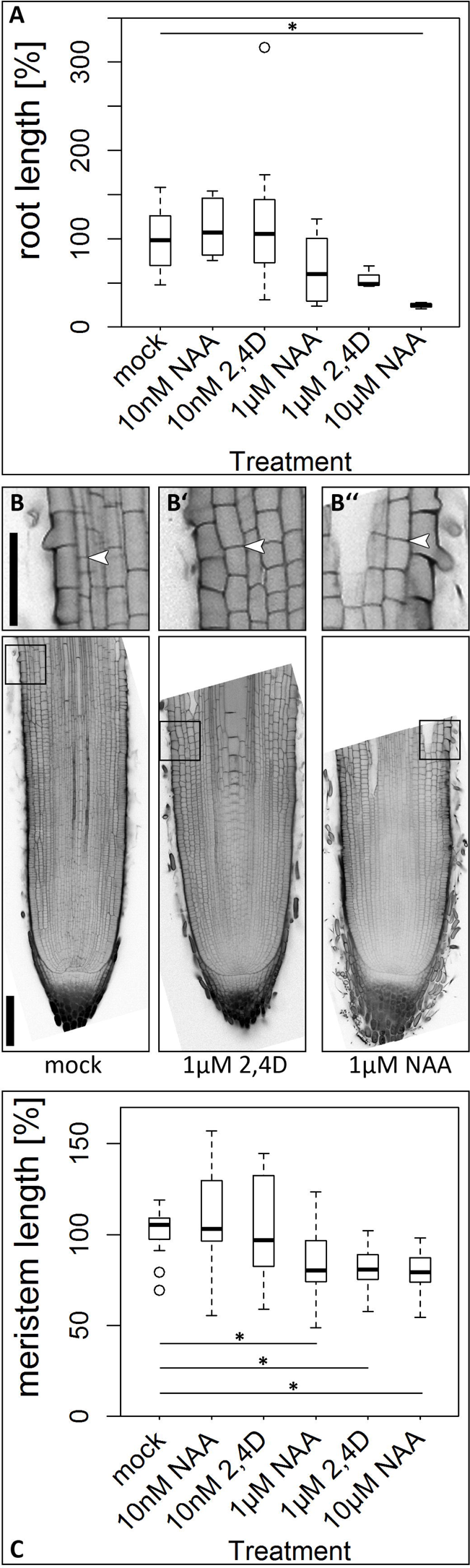
Root length, meristem size and DSC phenotype of the cv. Morex upon auxin treatment for 10 days. **A)** Root length after 10 day-treatment with auxin; experiment was performed twice; for a better comparison between the experiments, all values were normalized to the mock-treated plants; n = 4-18 plants per data point. **B) - B’’)** Representative pictures of the root meristem phenotype at 10 DAG upon hormone treatment according to the captions; arrow heads mark the transition zones; insets show magnifications of the transition zones; scale bars 200 μm and 100 μm in the magnification. **C)** Meristem length upon hormone treatment, measured by meristem length; experiment was performed twice; all values are normalized to the mock-treated control; n = 7-17 roots per data point; significance was determined using the two-tailed Student’s t test, * = p<0.05, **= p<0.001.

In *Arabidopsis*, auxin application results in differentiation of the DSCs, indicated by their accumulation of starch granules (Ding and Friml, 2010). In barley, no significant difference in the number of DSC layers was detected after auxin application (Supplementary figure 4).

### The commonly used auxin signalling reporter *DR5* and *DR5v2* are not stably expressed in barley

As auxin reporters, two widely used regulatory sequences are the *DR5* and the *DR5v2*, the former consisting of 9 inverted repeats of the auxin responsive element TGTCTC (Ulmasov et al., 1997) and the latter of 9 repeats of the higher affinity ARF binding-site TGTCGG (Liao et al., 2015). Here, the presence of auxin in a cell is indirectly determined through the activation of ARFs that bind to the synthetic promoters in an auxin-dependent manner, activating expression of the reporter genes. In *Arabidopsis*, the responsiveness of these reporters to auxin was confirmed by auxin application to the roots, leading to an enhanced expression of the reporter gene and a broadening of the expression domain (Liao et al., 2015). The same reporters were successfully used in rice and maize to display the spatial domain of auxin signalling (Yang et al., 2017; Gallavotti et al., 2008), therefore, the same regulatory sequences might be usable to monitor auxin signalling in barley. Surprisingly, no expression of the *DR5:GFP* reporter could be detected in transgenic lines with this construct, and expression of the *DR5v2:VENUS-H2B* reporter lines was very weak and inconsistent between different roots and plant lines (Supplementary figure 5A, B). Furthermore, no increase of *DR5v2:VENUS-H2B* expression was detected even upon high auxin concentrations (10 μM 2,4D) (Supplementary figure 5B’). Thus, the *DR5* and *DR5v2* reporters are not suitable to report on auxin signalling in barley.

### Expression pattern of *HvPLT1*

We therefore identified further genes known to be involved in auxin signalling, among them the PLT transcription factors (Aida et al., 2004; Galinha et al., 2007; Li and Xue, 2011). In *Arabidopsis*, the expression of the *AtPLTs* is dependent on auxin signaling, and the expression of AtPLT1 and AtPLT2 spatially reflects the auxin distribution in the root tip, observed by the expression of the DR5 auxin reporters (Mähönen et al., 2014; Galinha et al., 2007; Aida et al., 2004). We searched the barley proteome (Mayer et al., 2012) and created an unrooted tree of PLT family proteins from barley, rice, *Arabidopsis* and maize (Supplementary figure 6). The rice OsPLT1 protein grouped together with AtPLT1-3 and AtBBM (AtPLT4) and therefore might have a similar function in the stem cell niche maintenance (Li and Xue, 2011). Consequently, we focused on MLOC_76811 as the closest homologue of OsPLT1 (Supplementary figure 6), which we named HvPLT1 accordingly. *HvPLT1* consists of two repeats of the conserved AP2 DNA binding domain and a conserved linker region (Figure 4A, http://pgsb.helmholtz-muenchen.de/plant/barley/) similar to *AtPLT1* and *AtPLT2* from *Arabidopsis*. We created transgenic reporter lines that expressed *HvPLT1* fused to *mVENUS* under the control of 1929 bp of putative *HvPLT1* promoter sequences. The reporter lines showed expression of HvPLT1 with the maximum in the QC and the surrounding cells, which gradually decreased towards the root cap, the proximal meristems and the outer root layers (Figure 4B, B’). Non-transgenic control plants did not show any expression (Supplementary figure 8A, A’). RNA *in situ* hybridisations with a probe for *HvPLT1* confirmed this expression pattern (Figure 4C).

**Figure 4:**
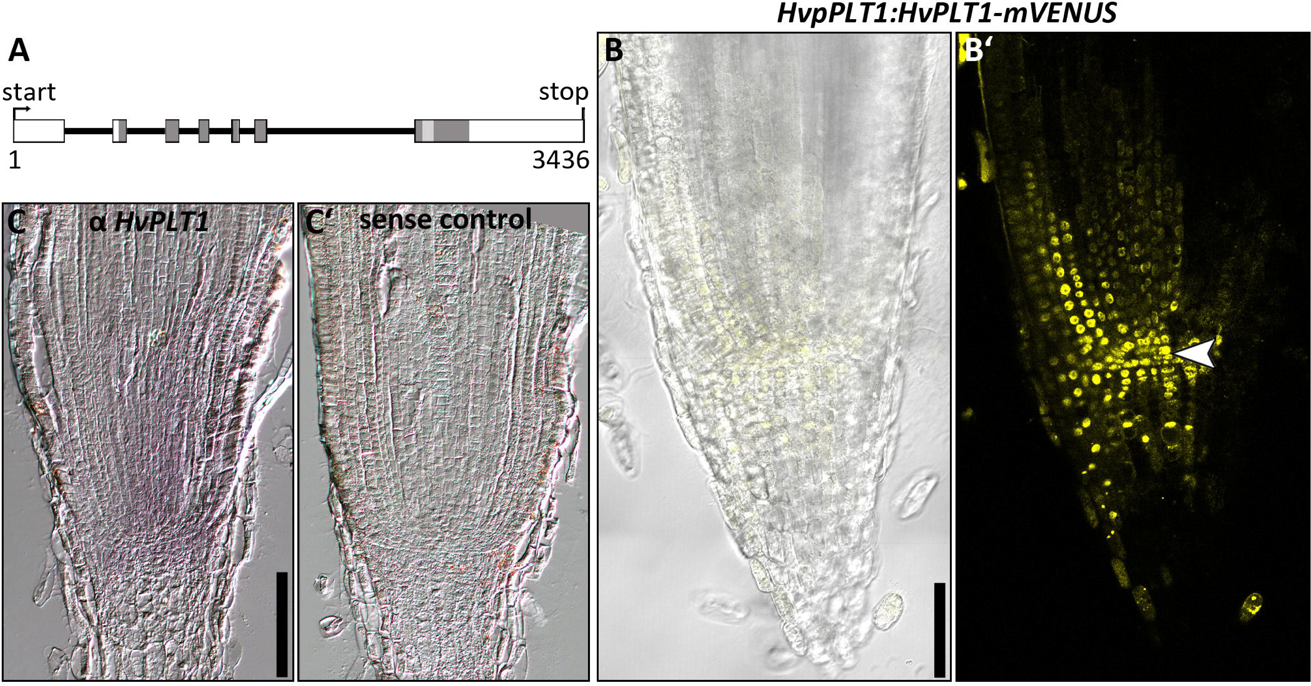
*HvPLT1* gene structure, promoter activity and protein localization in the root meristem of the barley cv. Golden Promise 8 DAG. **A)** Genomic structure of the *HvPLT1* coding sequence; boxes represent exons, black horizontal lines represent introns; dark gray boxes indicate coding sequence for AP2 domains, light gray boxes indicate coding sequence for the linkers between AP2 domains. **B)** Representative picture of the *HvpPLT1:HvPLT1-mVENUS* emission in the root meristem; transmitted light and mVENUS emission (B)), mVENUS emission only (B’)); arrow head in B’) points to the QC; hand sections; seven independent transgenic lines were examined and exhibited similar expression patterns. **C)** Representative picture of RNA *in situ* hybridizations with a probe for *HvPLT1* (purple staining, C)) or the respective sense probe (C’)); scale bars 100 μm.

### Identification of a PIN1 homologue in barley

The *HvPLT1* expression pattern suggests the presence of an auxin maximum in the QC and the root stem cell niche also in barley. A major source of auxin are young aerial tissues, from where the hormone is transported towards the root via the phloem (Saini et al., 2013) and subsequent cell-to-cell transport is facilitated by PIN proteins (Wang et al., 2009a; Carraro et al., 2006; Blilou et al., 2005). A search in the barley protein database for homologues of AtPINs discovered 13 putative HvPIN protein sequences that we used to build a phylogenetic tree and analyze their topology (Mayer et al., 2012) (Supplementary figure 9A). Based on the structure of the phylogenetic tree, we identified PIN1 (MLOC_64867.2, MLOC_12686.1), PIN2 (AK366549), PIN5 (MLOC_60446.1, MLOC_71135.1), PIN8 (MLOC_61956.2), PIN9 (MLOC_38112.1, MLOC_53867.1), PIN3 (MLOC_6128.3, MLOC_38023.1) and PIN10 (MLOC_38022.1, MLOC_60432.1) homologues and named them accordingly (Supplementary figure 9A). Like maize and rice, also barley does not encode *PIN4* and *PIN7* homologues (Wang et al., 2009a; Forestan et al., 2012). For maize it was hypothesized that in the root apex, the three ZmPIN1s could take over the roles of PIN3, PIN4 and PIN7 efflux carriers (Forestan et al., 2012). The same distribution of functions could hold true for barley. Additionally, the barley genome encodes one member of the SISTER OF PIN1 (SoPIN1) clade (MLOC_293.2, Supplementary figure 9A), which is conserved in flowering plants, but lost in *Arabidopsis* (O’Connor et al., 2014). A transmembrane helices prediction analysis revealed that the HvPIN1 and HvPIN2 carry 4 - 5 transmembrane domains that group around a central hydrophilic region, similar to AtPIN1 (Supplementary figure 9B). HvPIN5, HvPIN8 and HvPIN9 in contrast exhibit only a short central hydrophilic region (Supplementary figure 9). HvPIN3s and HvPIN10s did not show any typical structure of PIN proteins, and either lack a large hydrophilic region (HvPIN10a, HvPIN10b and HvPIN3b) or their hydrophilic region is not central (HvPIN3a) (Supplementary figure 9B). For the subsequent work on PIN protein localization in barley, we focussed on PIN1, as this is the best studied PIN protein in other model plants. Both in maize and rice, the two maize PIN1-like proteins and OsPIN1 show a similar transmembrane helices prediction profile, with two hydrophobic domains at the N and C termini and a central hydrophilic region (Xu et al., 2005; Wang et al., 2009a; Carraro et al., 2006). From the HvPINs that grouped together with the other PIN1s, HvPIN1a is the one with the transmembrane helices prediction profile most similar to AtPIN1 (Supplementary figure 9).

### Expression pattern and polar localization of HvPIN1a

In the *Arabidopsis* root, *PIN1* is expressed in the root meristem, in particular in the vasculature and endodermis, and weaker in the epidermis and cortex (Blilou et al., 2005). *PIN1* homologues of maize and rice are expressed in the root meristem, and also in the root cap (Forestan et al., 2012; Wang et al., 2009b). We generated transgenic barley reporter lines using the genomic *HvPIN1a* sequence under control of the putative endogenous *HvPIN1a* regulatory sequences, consisting of 3453 bp upstream of the start codon. The sequence of the fluorophore *mVENUS* was inserted into the part of the *HvPIN1a* gene sequence that encodes the intracellular hydrophilic region of the protein, as it was described for the AtPIN1-GFP reporter (Benková et al., 2003) (Supplementary figure 9B). Strong expression was detected in the whole root meristem, except for the area of the presumed QC, where expression was weaker compared to surrounding tissues (Figure 5A’, D’). High expression was observed in the stele, the endodermis, the cortex and DSCs, and the differentiated root cap (Figure 5D’). In the two analysed stages of the SAM, HvPIN1a was expressed throughout the whole meristem (Figure 6A, C). The PIN1s from *Arabidopsis*, maize and rice were shown to be mostly polarly localised to the plasma membranes at defined sides of the cells. In barley, a basal plasma membrane localisation was detected in the stele, endodermis and the inner cortex cell layers (gray arrow heads in Figure 5B, D’), but apical localisation was observed in the outermost cortex cell layer and the lateral root cap (white arrow head in Figure 5B, D’). In the central region of the root cap, polar localisation was not detectable, and HvPIN1a appeared evenly distributed at the plasma membrane (Figure 5D’). In the SAM, HvPIN1a was expressed everywhere, with expression peaks at the tips of the developing primordia (Figure 6). Here, we did not observe any polar localization of HvPIN1a (Figure 6).

**Figure 5:**
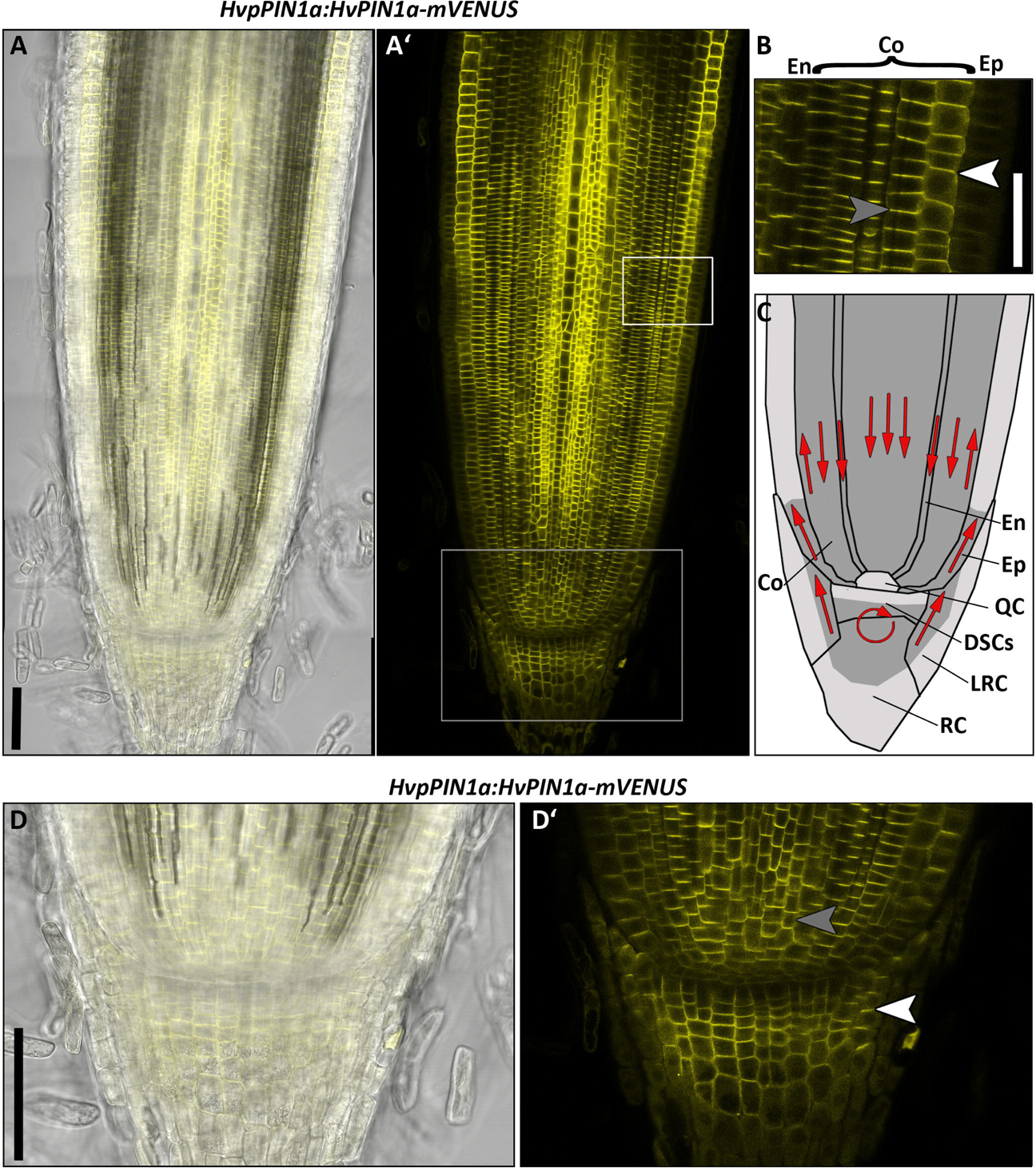
HvPIN1a expression in the root meristem of the barley cv. Golden Promise 8 DAG. **A)** Representative picture of *HvpPIN1a:HvPIN1a-mVENUS* expression; transmitted light and mVENUS emission (A)), mVENUS emission only (A’)); six independent transgenic lines were examined which vary only in expression level but not in localisation or pattern; white box in A’) marks magnification in B); gray box in A’) marks magnification in D). **B)** Magnification of the epidermal, cortical and endodermal cell layers depicted with white frame in A’). **C)** Schematic illustration of HvPIN1a expression in the root meristem, high = dark gray, low = gray; red arrows indicate possible auxin flow created by localisation of PIN1 auxin transporters; En = endodermis, Co = cortex, Ep = epidermis, LRC = lateral root cap, RC = root cap. **D)** Magnification of the stem cell niche depicted with gray frame in A’); transmitted light and mVENUS emission (D)), mVENUS emission only (D’)); white arrow heads mark apically localised PIN1, gray arrow heads mark basally localised PIN1; scale bars 100 μm in A), D); 50 μm in B).

**Figure 6:**
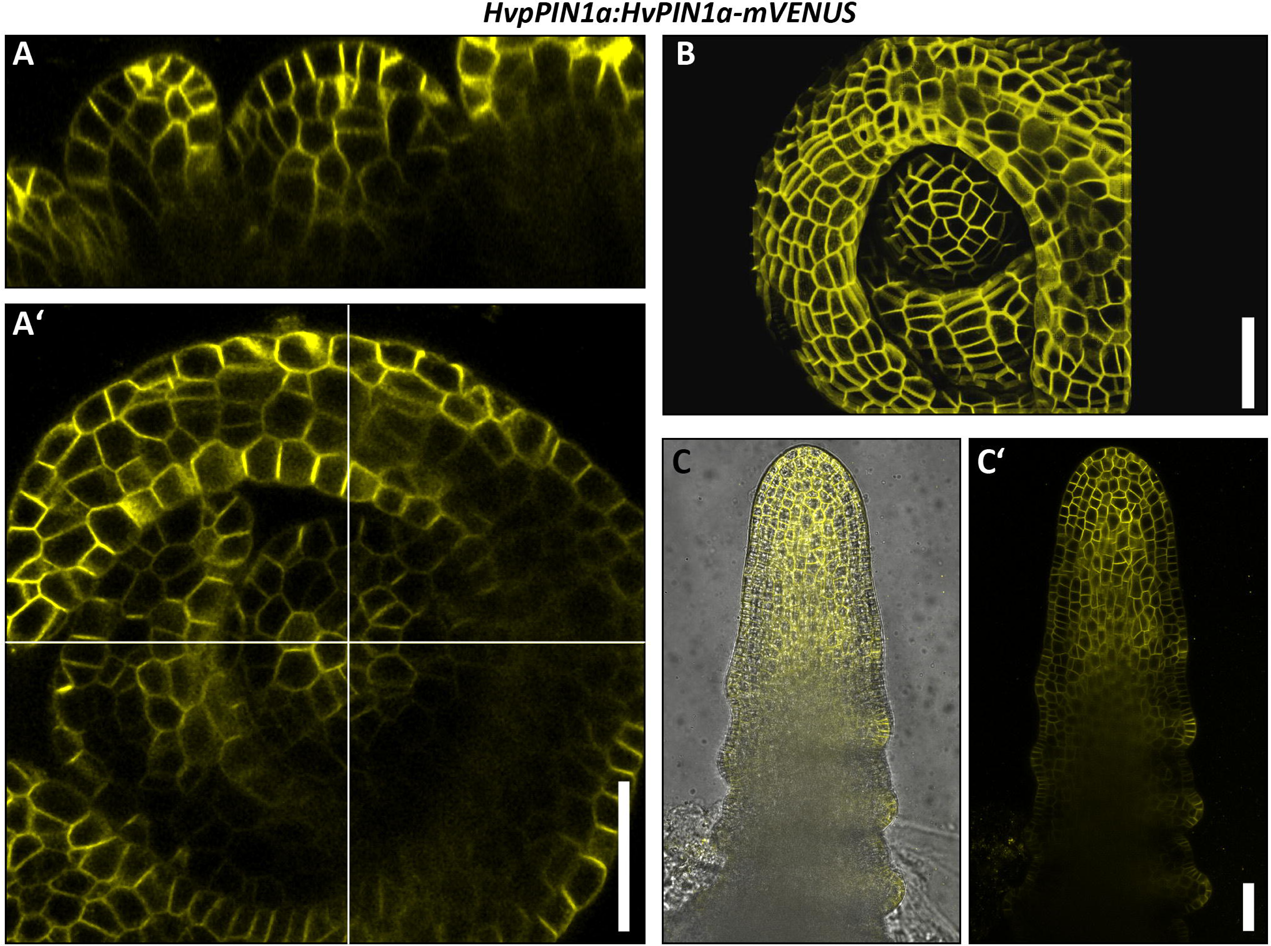
HvPIN1a expression in the shoot meristem of the barley cv. Golden Promise. **A), A’)** Representative picture of *HvpPIN1a:HvPIN1a-mVENUS* expression in the barley SAM in Waddington stage I; longitudinal view (A’)) and top view (A’)) of the same SAM. **B)** Surface projection of *HvpPIN1a:HvPIN1a-mVENUS* of the same SAM as in A), created with MorphoGraphX (de Reuille et al., 2015); **C)**, **C’)** Representative picture of *HvpPIN1a:HvPIN1a-mVENUS* expression in the barley SAM in Waddington stage II; transmitted light and mVENUS emission (C)), mVENUS emission only (C’)); four independent transgenic lines were examined and vary only in expression strength but not in localisation and pattern; scale bars 50 μm.

### HvPIN1a-mVENUS accumulates in vesicles upon Brefeldin-A (BFA) treatment

PINs are continuously recycled from the cell membrane to endosomes. Trafficking of basally localised PINs in *Arabidopsis* requires the GDP/GTP exchange factor for small G proteins of the ADP-ribosylation factor class (ARF-GEFs), which contain Sec7 domains (Kleine-Vehn et al., 2009; Steinmann et al., 1999; Geldner et al., 2003). The inhibitor of protein secretion BFA stabilizes an intermediate of the reaction of the ARF-GEF Sec7 domain with GDP, thereby blocking the cycle of activation of the ARF-GEFs and the recycling pathways (Peyroche et al., 1999). Therefore, BFA can be used to reveal the involvement of BFA-sensitive ARF-GEFs in the PIN recycling pathways. Treatment with BFA induces intracellular accumulation of AtPIN1 by blocking the exocytosis of PIN1, which normally cycles rapidly between plasma membrane and endosomal compartments (Geldner et al., 2001, 2003). To test if the PIN1 recycling mechanism is conserved in barley, we treated HvPIN1a-mVENUS expressing roots and SAMs with 50 μM BFA and monitored the HvPIN1a-mVENUS localisation after 2 h in the outer cortex cell file and the epidermis in root and shoot meristem, respectively. While HvPIN1a-mVENUS was exclusively localised at the apical cell membranes before the BFA treatment and upon mock controls, the formation of vesicles within the cells could be observed after 2 h of BFA treatment (gray arrow heads in Figure 7A). This indicates the existence of a conserved mechanism of PIN1 recycling between endosomal compartments and the plasma membrane in barley.

**Figure 7:**
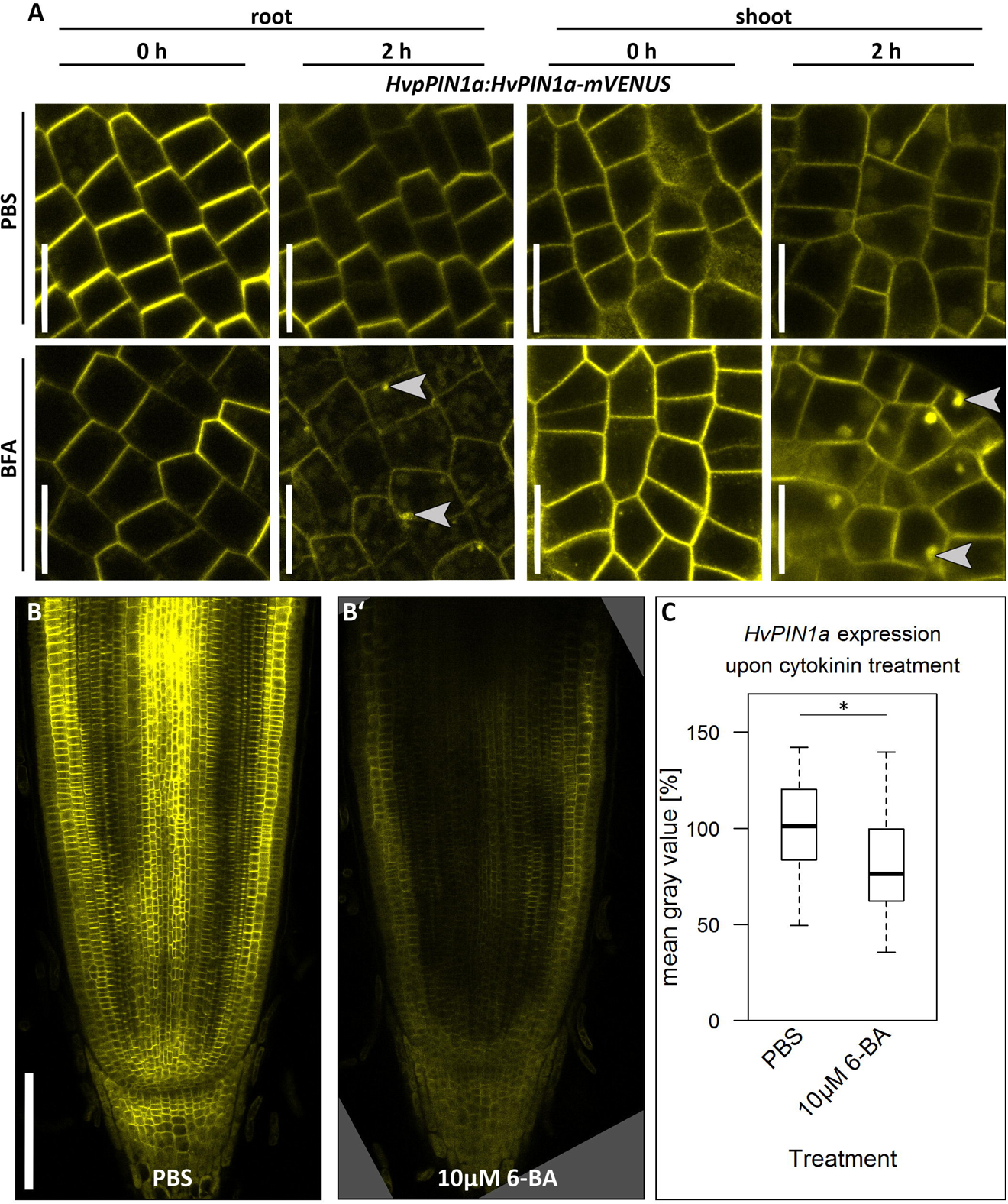
HvPIN1a localisation is influenced by BFA and its expression is decreased by cytokinin. **A)** Representative pictures of the *HvpPIN1a:HvPIN1a-mVENUS* expression in the outer cortex cell layer of the root meristem or in the epidermis of the SAM immediately after (0 h) or 2 h after mock (PBS) or 50 μM BFA treatment; gray arrow heads point to vesicles; scale bars 20 μm; three independent transgenic lines were examined; experiments were performed twice; n = 4 - 6; one transgenic line was used (in case of the root meristem); experiments were performed twice; n = 3 - 5; two transgenic lines were used (in case of shoot meristem). **B)** Representative pictures of *HvpPIN1a:HvPIN1a-mVENUS* expression upon either mock (PBS, B)) or cytokinin (B’)) treatment as indicated in the captions; scale bar 200 μm. **C)** Quantitative analysis of the *HvpPIN1a:HvPIN1a-mVENUS* expression upon cytokinin expression in B), measured by the mean gray value of the whole root meristem and the root cap; values are normalized to the PBS-control; five different independent transgenic lines were used; experiment was performed twice; n = 24 per treatment; significance was determined using the two-tailed Student’s t test, * = p<0.05.

### *HvPIN1a* expression is regulated by cytokinin

As the recycling of the HvPIN1a protein is similarly affected by BFA as it is in *Arabidopsis*, we examined if gene expression of *HvPIN1a* is regulated by the same factors as in *Arabidopsis*. Dello Ioio and colleagues showed that *AtPIN1* expression is downregulated by cytokinin (Dello Ioio et al., 2008; Ruzicka et al., 2009). In barley, treatment with the cytokinin 6-BA for 24 h reduced *HvPIN1a-mVENUS* expression as well (Figure 7B, C). The downregulation was mostly visible in the stele, the place of cytokinin signaling (Figure 7B). However, if this downregulation occurs at the level of transcription like in *Arabidopsis*, or on protein level, we cannot distinguish here (Dello Ioio et al., 2008).

## Discussion

### Creation and validation of auxin and cytokinin reporter lines in barley

With this study, we started to address the roles of auxin and cytokinin in barley root and shoot meristem development at cellular resolution. We found that the *DR5* and *DR5v2* regulatory elements that are successfully used in many other plant species to determine auxin signalling were not consistently expressed in our transgenic barley lines (Liao et al., 2015; Yang et al., 2017; Forestan et al., 2012) (Supplementary figure 5). In *Arabidopsis*, the two auxin reporters *DR5* and *DR5v2* showed differences in their expression patterns, indicating that ARFs have different binding affinity towards the respective auxin response elements TGTCTC (*DR5*) and TGTCGG (*DR5v2*) (Liao et al., 2015). Moreover, it was shown that the spacing between the auxin responsive elements, their flanking sequences and the number of repeats are important for the reactivity of the reporter to auxin (Ulmasov et al., 1997). We have to conclude from the data presented here that in barley, different auxin responsive elements and/or a different composition of the reporters are necessary to create transcriptional reporter lines for auxin signalling. We could, however, successfully monitor the expression of auxin-related genes. We found that the expression pattern of HvPLT1 in root meristems is similar to that of AtPLT1 in *Arabidopsis* and the root-specific rice PLTs, with an expression maximum around the QC (Galinha et al., 2007; Li and Xue, 2011), both on RNA and protein level (Figure 4). This suggests that the expression of *PLTs* is conserved between these plant species, indicating that the well described auxin- and PLT-mediated cell specification mechanism in the root meristem is conserved between *Arabidopsis* and barley (Aida et al., 2004; Galinha et al., 2007; Mähönen et al., 2014). The expression of PIN1 in the root is conserved between species, but expression in individual tissues differs (Figure 5A’) (Wang et al., 2009a; Blilou et al., 2005; Forestan et al., 2012). Furthermore, the intracellular localisation of PINs differs depending on their domain topology. Křeček and colleagues sorted the eight *Arabidopsis* PINs into two subfamilies, namely the “long” and the “short” PINs according to the length of their hydrophilic region (Křeček et al., 2009). The “long” PIN subfamily is characterised by its central hydrophilic loop, separating two hydrophilic domains, each consisting of five trans-membrane regions. They are primarily localised to the plasma membrane in the cell (Benková et al., 2003; Blilou et al., 2005; Friml et al., 2003). The “short” PINs, however, possess a short central hydrophilic region and localise to internal cell membranes (Ganguly et al., 2010). Accordingly, HvPIN1a localised to the plasma membrane (Figure 5B, D’). We furthermore discovered PINs that cannot be assigned as “long” or “short” PIN because their transmembrane topology does not follow either structure (Supplementary figure 9B). This divergence in transmembrane topology has not been reported for PINs in *Arabidopsis*, rice and maize and therefore, the localisation and function of these PINs should be subjected to a closer examination. The expression pattern and polar localization of HvPIN1a indicate that also in barley, an auxin flow is created that is directed towards the QC, the stem cell niche and the root cap and also a flow from the stem cell niche to the proximal meristem via the outer cortex cell layers (Figure 5C), as it was proposed for the *Arabidopsis* PINs (Blilou et al., 2005). In the SAM, AtPIN1 is expressed in the epidermis and in subepidermal cells, and only below the primordia, the expression becomes restricted to the presumed provascular tissues (Heisler et al., 2005). In maize, ZmPIN1a is also predominantly expressed in the epidermis of axillary meristems and the inflorescence meristem, but also in underlying tissues (Gallavotti et al., 2008). Like in the root, HvPIN1a is more broadly expressed than PIN1 in *Arabidopsis*, not only in the epidermis but evenly in all tissue layers of the SAM (Figure 6). Like in the root, it is possible that close PIN1 homologues take over the role of a tissue-layer specific auxin transporter in barley shoots (Supplementary figure 9A).

Genetic analysis and BFA-treatment experiments revealed that in *Arabidopsis*, the basal localisation of PINs is dependent on the ARF-GEF GNOM (Geldner et al., 2003). The kinases D6 PROTEIN KINASE and PINOID regulate PIN localization by phosphorylating the PINs at the plasma membrane, making them less affine to the GNOM-dependent basal recycling pathway. The phosphorylated PIN proteins are then recruited to the apical GNOM-independent trafficking pathways (Kleine-Vehn et al., 2009; Steinmann et al., 1999; Geldner et al., 2003). For the apically localised AtPIN2 in the *Arabidopsis* epidermis, however, it was shown that its vacuolar trafficking is independent of GNOM and involves an additional, BFA-sensitive ARF-GEF (Kleine-Vehn et al., 2008). In the outer cortex cell layer, where HvPIN1a is localised apically, and in the SAM epidermis, where we could not observe a polar localisation, BFA caused the accumulation of HvPIN1a in vesicles (Figure 7A), indicating that in barley, too, BFA-sensitive components are involved in PIN1 trafficking. Besides intracellular localisation of the PIN1 protein, also the expression of *PIN1* is subject to regulation by other factors. Dello Ioio and colleagues showed that in *Arabidopsis*, *AtPIN1* expression is downregulated by cytokinin (Dello Ioio et al., 2008). In rice, the PIN1 homologues *OsPIN1a*, *OsPIN1b* and *OsPIN1c*, however, are not transcriptionally regulated by cytokinin (Wang et al., 2009b). In barley, *HvPIN1a* expression is downregulated by cytokinin application (Figure 7B, C). Thus, *HvPIN1a* expression is similarly regulated in barley as it is in *Arabidopsis*, but different from rice, where *OsPIN1* expression is cytokinin independent (Dello Ioio et al., 2008; Wang et al., 2009b).

In contrast to the auxin reporters *DR5* and *DR5v2*, the cytokinin signaling sensor *TCSn* was functional in barley, since its expression pattern is conserved between Arabidopsis and barley, and, more importantly, the expression pattern changes upon cytokinin application (Figure 2) (Zurcher et al., 2013). However, although the *TCSn* reporter confers expression in the shoot meristem in *Arabidopsis* (Zurcher et al., 2013), we could not detect any expression of *TCSn:VENUS-H2B* in barley SAMs. Since the *TCSn* reporter consists of concatemeric repeats of the DNA-binding motif of the *Arabidopsis* type-B ARR (Zurcher et al., 2013), it is possible that in barley, only some tissues express proteins that induce the *TCSn* regulatory sequence, so that despite ongoing cytokinin signalling in the cell, the reporter gene is not activated.

### Role of auxin and cytokinin for barley root meristems

Previous studies on hormone activities in barley had addressed how manipulation of endogenous cytokinin levels and auxin affects root growth, at the whole organ level (Zalewski et al., 2010; Pospíšilová et al., 2016; Tagliani et al., 1986). We have extended these analyses to the meristem and cellular level. The reduction in root growth as well as a reduction in meristem size upon 6-BA application in barley is similar to that observed for *Arabidopsis* (Figure 1) (Ruzicka et al., 2009; Dello Ioio et al., 2007). In *Arabidopsis*, components of cytokinin signalling in the root stem cell niche control the differentiation at the transition zone (Dello Ioio et al., 2007; Moubayidin et al., 2013). Expression of the cytokinin reporter *TCSn:VENUS-H2B* in barley was enhanced in the stem cell niche upon cytokinin treatment, indicating that the reduction of meristem size upon cytokinin treatment might also depend on enhanced cytokinin signalling in the QC, as observed for *Arabidopsis* (Moubayidin et al., 2013). Interestingly, all effects on root growth and root meristem size were less pronounced upon treatment with the cytokinin t-Z compared to 6-BA. In *Arabidopsis*, CYTOKININ OXIDASES/DEHYDROGENASES (AtCKXs), which are involved in the degradation of cytokinins, preferentially cleave isoprenoid cytokinins including t-Z, but not 6-BA; similarly, CKX1 from maize predominately cleaves free cytokinin bases including t-Z (Galuszka et al., 2007; Mrízová et al., 2013). In barley, thirteen putative members of the *HvCKX* family were identified (Zalewski et al., 2014). Their presence could lead to an enhanced degradation of the externally added t-Z, thereby leading to a reduced influence on root growth and meristem maintenance in comparison to 6-BA.

Besides cytokinin, we analysed the influence of the synthetic auxin NAA and the non-transportable synthetic auxin analogue 2,4D on barley root growth. In contrast to studies in *Arabidopsis* where low concentrations of auxins were shown to increase the root growth rate (Evans et al., 1994; Müssig et al., 2003), we observed no enhancement of root growth rates upon applications of low concentrations of NAA and 2,4D (10 nM) in barley (B). Instead, we observed a reduction in meristem size upon treatment with high concentrations of auxin, both in regard to cell number and meristem length (Figure 3, Supplementary figure 1). Besides the effect of auxin application on longitudinal root growth and meristem size in *Arabidopsis*, the phytohormone also influences the DSCs that give rise to the columella cells. Auxin application leads to a differentiation of these stem cells, marked by accumulation of starch granules (Ding and Friml, 2010). In barley, however, we could not observe any starch granule accumulations in additional cell files (Supplementary figure 4B). Previously, we have published similar observations for the application of a CLE peptide. CLE peptides were shown to cause both a differentiation of the proximal root meristem and the DSCs in *Arabidopsis*. Application of the CLE peptide in barley, however, did only affect the proximal root meristem but not the DSC differentiation (Kirschner et al., 2017; Stahl et al., 2009). This indicates that DSC maintenance, in contrast to root meristem maintenance, is regulated differently in barley than in *Arabidopsis*.

In summary, we have shown here important similiarities and differences in root meristem development and the role of phytohormones between barley, other crop species and the model organism Arabidopsis. We have also characterised a first set of fluorescent reporter lines for barley at cellular resolution, which will be useful for further in-depth studies of the poorly understood development of one of the major crop plants worldwide.

## Supplementary material

**Supplementary figure 1: Meristem cell number upon auxin and cytokinin treatment A)** Meristem cell number upon 10-day cytokinin treatment; experiment was performed twice; n = 11-16 roots per data point. **B)** Meristem length upon auxin treatment; experiment was performed twice; all values are normalized to the mock-treated control; n = 7-17 roots per data point; significance was determined using the two-tailed Student’s t test, * = p<0.05, **= p<0.001.

**Supplementary figure 2: Root meristem width upon cytokinin and auxin treatment.** Meristem width measured at the transition zone from root meristems exemplarily shown in Figure 1B and Figure 3B. **A)** Roots were treated with cytokinin for 10 days; experiments were performed twice; n = 15-25 roots per data point. **B)** Roots were treated with auxin for 10 days; experiment was performed twice, n = 12-25 roots per data point; significance was determined using the two-tailed Student’s t test, * = p<0.05, **= p<0.001.

**Supplementary figure 3: The cytokinin reporter *TCSn:VENUS-H2B* is not expressed in the SAM of the barley cv. Golden Promise. A), A’)** Undectectable *TCSn:VENUS-H2B* expression in SAMs in waddington stage I; transmitted light and VENUS emission (A)) and VENUS emission only (A’)). **B), B’)** Undectectable *TCSn:VENUS-H2B* expression in SAMs in waddington stage II; transmitted light and VENUS emission (B)) and VENUS emission only (B’)). Seven independent transgenic lines were examined and show no expression in the SAM; scale bars 100 μm; insets in A’) and B’) show respective pictures with tonal correction to show autofluorescence.

**Supplementary figure 4: DSC layer number of the cv. Morex upon 10-day treatment with auxin. A)** Exemplary pictures of the root stem cell niche upon mock or auxin treatment as indicated; scale bar 100 μm. **B)** Number of DSC layers upon 10-day treatment with auxin; no significant difference to mock-treated plants; experiment was performed twice; n = 5-20 per data point. Significance was determined using the two-tailed Student’s t test, * = p<0.05, **= p<0.001.

**Supplementary figure 5: Expression of *DR5* is undetectable and *DR5v2* is unstably expressed in barley roots. A)** Exemplary picture of *DR5:GFP* root; transmitted light and (undetectable) GFP emission (A)), undetectable GFP emission only (A’)); four independent transgenic lines were examined and show no GFP epression; scale bar 200 μm. **B)** *DR5v2:VENUS-H2B* lines show only variable or no expression and do not show a consistent reaction on 2,4D treatment; number of expressing *DR5v2:VENUS-H2B* lines (B)); number of plants that show the respective expression change upon treatment with 10 μM 2, 4D for 24 h (B’)).

**Supplementary figure 6: Phylogenetic tree of PLT homologue proteins.** *Arabidopsis* PLT sequences were taken from arabidopsis.org; rice PLT sequences were named according to Li and Xue (Li and Xue, 2011); maize PLT sequences were identified in a BLAST search with AtPLT1 as template (e-value below 5e-75) on the Phytozomev.12.0 website and named according to Zhang and colleagues (Zhang et al., 2014); barley genes were identified by BLAST-p search on http://webblast.ipk-gatersleben.de/barley/ with AtPLT1 as template (e-value below 4e-47 for high-confidence genes and 2e-11 for low-confidence genes) (Mayer et al., 2012). Alignments and evolutionary analyses were performed using MEGA7. 0 (Molecular Evolutionary Genetics Analysis version 7.0 for bigger datasets (Kumar, Stecher and Tamura 2015)) and a MUSCLE alignment; the evolutionary history was inferred by using the Maximum Likelihood method based on the JTT matrix-based model. The bootstrap consensus tree inferred from 100 replicates is taken to represent the evolutionary history of the taxa analyzed. Branches corresponding to partitions reproduced in less than 50% bootstrap replicates are collapsed. The percentage of replicate trees in which the associated taxa clustered together in the bootstrap test (100 replicates) are shown next to the branches. Initial tree(s) for the heuristic search were obtained automatically by applying Neighbor-Join and BioNJ algorithms to a matrix of pairwise distances estimated using a JTT model, and then selecting the topology with superior log likelihood value.

**Supplementary figure 7: Barley cv. Golden Promise as non-transgenic control. A)** Representative picture of the root meristem of a non-transgenic Golden Promise seedling 8 DAG; transmitted light and mVENUS emission (A)), mVENUS emission only (A’)), same settings as in Figure 4B, B’; hand-sections as described in Material and Methods; only background signal with mVENUS excitation. **B)** Representative picture of the root meristem of a non-transgenic Golden Promise seedling 8 DAG; transmitted light and mVENUS emission (B)), mVENUS emission only (B’)), same settings as in Figure 5A’; cleared as described in Material and Methods; only background signal with mVENUS excitation; scale bars 100 μm; inset in A’) shows respective pictures with tonal correction to show autofluorescence.

**Supplementary figure 8: Phylogeny and topology of barley PINs. A)** Phylogenetic tree of maize, *Arabidopsis*, rice and barley PINs; barley PINs were taken from http://webblast.ipk-gatersleben.de/barley/ with BLAST-p with HvPIN1a (MLOC_64867) as template (e-value below 1e-41 for high and low-confidence genes); rice sequences are taken from (Miyashita et al., 2010); Arabidopsis PINs were searched at arabidopsis.org; maize PINs were taken from Phytozome v12 (e-value below 4.3e-29) and named according to (Forestan et al., 2012). Selected SoPIN1 proteins from Brachypodium and tomato (Solanum lycopersicum) taken from (O’Connor et al., 2014; Martinez et al., 2016) to define the SoPIN1 clade. Alignments were performed using MEGA7. 0 (Molecular Evolutionary Genetics Analysis version 7.0 for bigger datasets (Kumar, Stecher and Tamura 2015)) and a MUSCLE alignment; the phylogenetic tree was obtained using MEGA7.0 by the Maximum Likelihood method based on the JTT matrix-based model. The bootstrap consensus tree inferred from 100 replicates is taken to represent the evolutionary history of the taxa analyzed. Branches corresponding to partitions reproduced in less than 50% bootstrap replicates are collapsed. The percentage of replicate trees in which the associated taxa clustered together in the bootstrap test (100 replicates) are shown next to the branches. Initial tree(s) for the heuristic search were obtained automatically by applying Neighbor-Join and BioNJ algorithms to a matrix of pairwise distances estimated using a JTT model, and then selecting the topology with superior log likelihood value; protein subfamilies are framed with the same colour; gray frame marks HvPIN1a. **B)** Topology of the transmembrane barley PIN proteins in comparison to AtPIN1; domains predicted to the inside of the cell are shown in light-gray, transmembrane domains are shown in dark-gray and domains outside the cell are depicted in black according to the legend; in the protein topology of MLOC_64867 - HvPIN1a the asterisk marks the site where mVENUS is inserted for the reporter line shown in Figure 5; newly identified HvPINs are named according to their topology and the cluster of the Arabidopsis, maize and rice PIN family to which they belong.

